# Taste dysfunction in Long COVID

**DOI:** 10.1101/2025.07.17.661973

**Authors:** Hanna Morad, Tytti Vanhala, Marta A. Kisiel, Agnes Andreason, Mei Li, Göran Andersson, Göran Laurell, Thomas E. Finger, Göran Hellekant

**Affiliations:** Clinic of Ear, Nose and Throat diseases (ENT), Uppsala University Hospital, Uppsala, Sweden; Department of Animal Biosciences, Swedish University of Agricultural Sciences, Uppsala, Sweden; Occupational and Environmental Medicine, Department of Medical Sciences, Uppsala University, Uppsala, Sweden; Rocky Mountain Taste & Smell Center, Department of Cell & Developmental Biology, University of Colorado School of Medicine, Aurora CO 80045, USA; Department of Surgical Sciences, Uppsala University, Uppsala, Sweden; University of Wisconsin Madison, School of Veterinary Medicine, Madison, WI, USA

**Author notes:** Shared Senior authors (Thomas E. Finger, Göran Hellekant). CORRESPONDING AUTHORS: Correspondence to Göran Hellekant.

## Abstract

Persistent taste dysfunction is frequently reported in individuals with post-acute sequelae of infection by SARS-CoV-2 (Long COVID). The mechanisms and pathological correlates underlying this taste dysfunction are unknown. This study investigates the underlying pathology in 28 non-hospitalized subjects diagnosed with COVID-19 and who experienced taste disturbances more than 12 months after testing positive for SARS-CoV-2. To objectively establish the nature of taste deficit, we used the WETT taste test, which quantifies the subject’s ability to taste each of the five taste qualities: sweet, umami, bitter, sour, and salty. We then biopsied five to eight fungiform taste papillae (FP) in 20 of the 28 subjects. The FPs were analyzed histologically for overall taste bud structure and innervation, and by quantitative PCR (qPCR) for mRNA expression of markers for different taste receptor cells. Although all subjects had reported taste dysfunction, only three showed overall taste scores below the 10^th^ percentile for a normal population adjusted for age and sex. However, 11 of the 28 subjects exhibited total loss of one or more taste qualities. Loss of PLCβ2-dependent taste qualities (sweet, umami, bitter) was significantly more common and was correlated with reduced expression of *PLCβ2* and *Tas1R3* mRNAs. Histological analysis revealed generally preserved taste bud structure and innervation, but with occasional disorganized taste buds and abnormal, isolated PLCβ2-positive cells in the epithelium. Our findings suggest long-term taste dysfunction after COVID-19 occurs rarely -- more frequently involving PLCβ2-dependent taste qualities -- but is not due to wholesale disruption of the taste periphery.

## INTRODUCTION

The COVID-19 pandemic has brought unprecedented attention to impairments of olfaction and taste as hallmark symptoms of acute Severe Acute Respiratory Syndrome-CoronaVirus-2 (SARS-CoV-2) infection (Cooper *et al*. 2020; Hannum *et al*. 2023; Klein *et al*. 2021a; Klein *et al*. 2021b). In most COVID-19-affected individuals, this dysfunction resolves within weeks but can persist for months for others. Prolonged dysfunction lasting more than three months is now recognized as part of post-acute COVID-19 syndrome (PACS) or Long COVID (Ely *et al*. 2024; Soriano *et al*. 2022). Ongoing taste dysfunction may occur even after resolution of other symptoms (Boldes *et al*. 2024; Xu *et al*. 2024) thus raising questions about the underlying mechanisms. Understanding the long-term pathophysiology of taste dysfunction and, in extreme cases, taste loss is crucial for patient care and the development of targeted rehabilitation strategies (Lee *et al*. 2025).

Sensations of taste are largely mediated by lingual taste buds (TBs) situated in three types of papillae: fungiform, vallate, and foliate. The fungiform papillae (FP) are concentrated on and around the tip of the tongue, making them easy to biopsy. A typical human has about 200 FP with 0-50 taste buds (TB) in each FP (Cheng and Robinson 1991; Miller 1986; Miller 1988 ; 1989 ; Miller and Reedy 1990).

Taste buds are complex multicellular epithelial end organs comprising between 20 and 100 taste cells (TCs) with approximately half being taste receptor cells (TRCs) (High and Finger 2025; Ogata and Ohtubo 2020; Ohtubo and Yoshii 2011; Yang *et al*. 2020). Histochemical, immuno-histochemical and molecular methods allow for correlation of TRC subtypes with specific taste qualities. Type II TRCs detect sweet, umami, or bitter tastes via distinct G-protein-coupled receptor (GPCR) pathways involving downstream PLCβ2-dependent signaling cascades (Shi *et al*. 2003; Zhang *et al*. 2003). Sweet taste is mediated via TAS1R2/R3 heterodimeric receptors, and umami by TAS1R1/R3; bitter taste involves TAS2Rs, another family of taste GPCRs. In contrast, Type III cells are responsible for transduction of ionic taste qualities, i.e. sour and high concentrations of salt (Chang *et al*. 2010; Matsumoto *et al*. 2011) while Type I TCs play a role similar to glia cells in the CNS (Bartel *et al*. 2006; Rodriguez *et al*. 2021; Wilson *et al*. 2025). In addition, taste buds contain a population of post-mitotic Type IV cells which mature to differentiate into the other cell types. Disruption to these distinct molecular cascades or to the morphological integrity of taste buds could contribute to the observed taste dysfunction in Long COVID (Huang *et al*. 2021a).

Taste bud cells undergo continuous turnover, with new TRCs differentiating only when taste fibers reach them (Farbman 1965; 1980; Farbman *et al*. 2004; Hellekant *et al*. 1987; Lee *et al*. 2017). Approximately 70–80% of mature TRCs form synapses with nerve fibers, highlighting the importance of neural connectivity in maintaining taste function (Lee *et al*. 2017; Wilson *et al*. 2025; Wilson *et al*. 2022). Ongoing post-viral disruption of the nerve-TC interactions could well lead to taste dysfunction.

SARS-CoV-2 infects cells via the ACE2 receptor and the TMPRSS2 protease, both of which are expressed in oral epithelial cells, including Type II TRCs (Doyle *et al*. 2021; Yao *et al*. 2023). This suggests a plausible mechanism for direct viral-caused damage to the taste system. Additionally, inflammatory processes, immune dysregulation, or altered saliva composition may contribute to long-term taste changes following viral infection.

Given the clinical and biological implications of long-term taste dysfunction, this study aims to investigate self-reported and objectively measured taste impairment in subjects who experienced non-hospitalized COVID-19. Using a combination of psychophysical testing, histological evaluation and molecular analysis -- including quantitative polymerase chain reaction (qPCR) for mRNAs encoding taste cell markers and viral RNA -– this study seeks to elucidate the structural and molecular changes underlying persistent taste dysfunction in Long COVID subjects following COVID-19 caused by SARS-CoV-2.

## MATERIAL & METHODS

### Study design

This study involved subjects aged ≥18 years who had experienced symptomatic COVID-19 and a confirmed SARS-CoV-2 infection verified by a positive PCR test from nasopharyngeal swabs between 2020 and 2021 (inclusion criteria). All subjects had non-hospitalized COVID-19 and, all but one, reported long-term taste disturbances. The study population was recruited between November 2021 and June 2022 from two sources at Uppsala University Hospital: the COMBAT post-COVID study and the Ear, Nose and Throat (ENT) outpatient clinic.

From the COMBAT post-COVID cohort, 18 subjects underwent psychophysical taste evaluation. Of these, 12 consented to undergo a fungiform papilla biopsy (subjects #SWE7, 8, 9, 12, 13, 14, 15, 18, 19, 23, 24, 26). Additionally, ten subjects were recruited from the ENT outpatient clinic, eight of whom underwent biopsy (subjects #SWE17, 21, 22, 24, 25, 27, 28, 29). In total, eight subjects did not undergo a biopsy.

### Study population

The COMBAT post-COVID project aimed to monitor persistent symptoms and long-term consequences among non-hospitalized patients diagnosed with COVID-19. Initially, 725 individuals who tested positive for SARS-CoV-2 between March and December 2020 were invited to complete a follow-up questionnaire 12 months after symptom onset. Of these, 401 subjects responded (response rate: 74%). The cohort has been described in detail elsewhere (Janols *et al*. 2024; Kisiel *et al*. 2022; Kisiel *et al*. 2023a; Kisiel *et al*. 2023b; Kisiel *et al*. 2021). Among the respondents, 68 individuals reported ongoing taste and/or smell disturbances. These individuals were contacted by telephone and invited to participate in the current study if they reported persistent taste loss and were available for in-person evaluation. Eighteen individuals agreed to participate. Henceforth all individuals will be indicated as subjects.

Ten additional subjects attending the ENT outpatient clinic in 2021 were randomly selected and invited to participate in the study if they met the same inclusion criteria. In addition, one subject (#SWE17) reported no taste loss but had an active SARS-CoV-2 infection at the time of sampling. Taste testing data for this subject are listed in Table 1 but are not included in the statistical analysis of the cohorts reporting Long COVID taste disturbance.

**Table 1:**
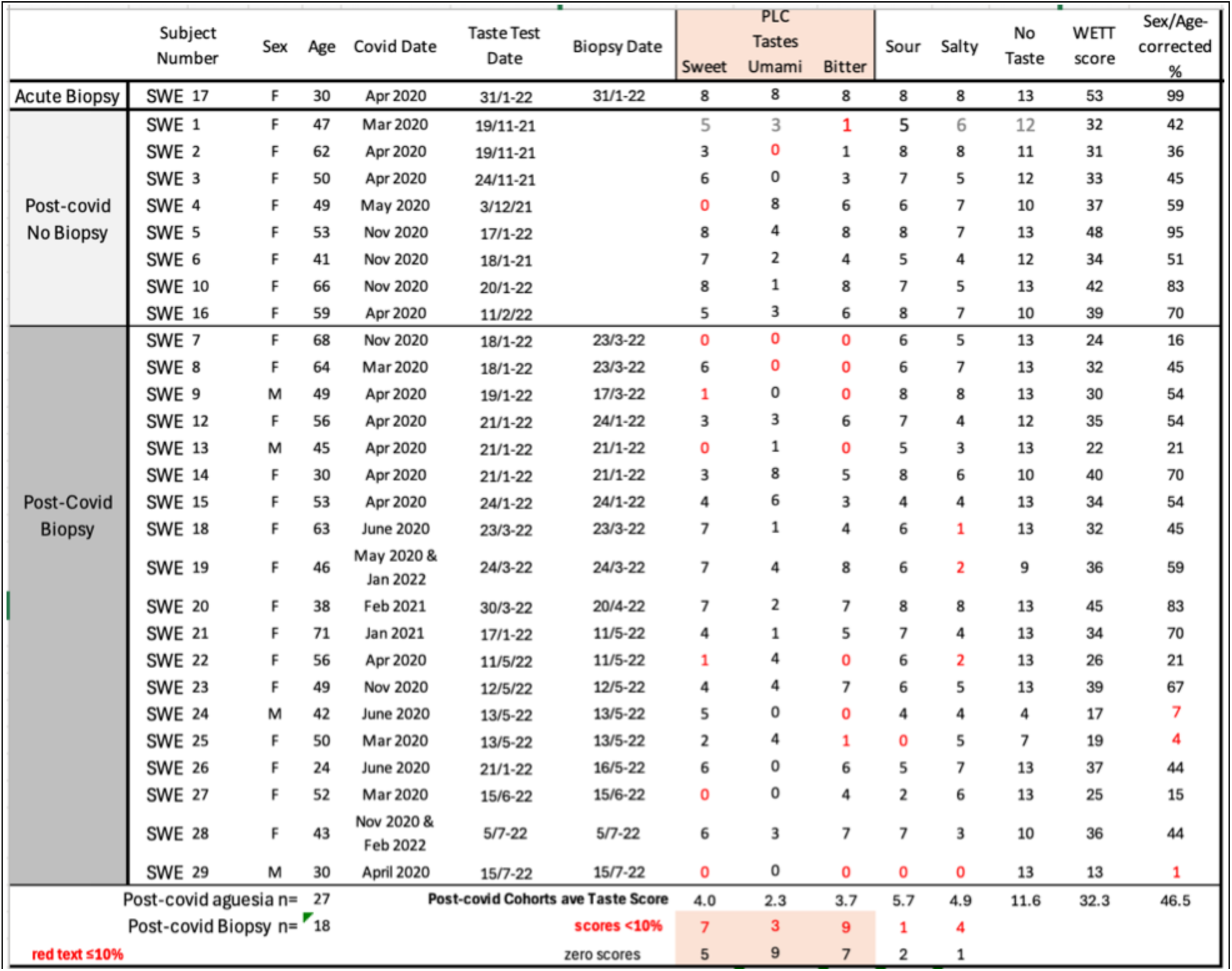
Results from WETT testing of all subjects including basic demographic information, dates of infection, taste testing and biopsy. Columns labeled as “Sweet, Umami, Bitter, Sour, Salty and No Taste” show WETT test raw scores for each taste quality and blank “No Taste” control strips. Red numerals in columns for individual tastes indicate values below the 95% confidence interval for sex and age-matched controls based on Doty, et al 2025. The incidence of values below the 10% for all post-Covid subjects is significantly higher than expected based on a normal population (χ²(1, N = 135) = 4.47, p < .05). Zero values indicate that the subject was unable to identify the tastant at any concentration offered. The total number of zero values for each tastant is shown in red along the bottom row. WETT score shows the total WETT score for each subject alongside of the corresponding sex and age matched percentile ranking compared to the study population in Doty et al 2025. A perfect score is 53 (8 for each taste quality + 13 blanks). Complete ageusia would give a score of 13 (0 for all tastants and 13 for correctly identifying blank controls). PLCß2-dependent tastes are indicated by orange shading along the top and bottom rows.

### Clinical and sociodemographic variables

From the COMBAT post-COVID cohort, information was documented about age, sex, symptoms at COVID-19 onset, self-reported health before and one year after COVID-19, and persistent symptoms one year following COVID-19. Symptoms at COVID-19 onset were also retrieved from medical records from the PCR testing protocols. Additional subjects (n=11) were characterized by age and sex only.

### Ethical Approval

The study was approved by the institutional ethics committee of the University Hospital in Uppsala (EPN number 2020-05707) and the Swedish Ethical Review Authority (EPM number 2021-00878) and complies with the Declaration of Helsinki for Medical Research involving Human Subjects. The patients all gave written informed consent to the procedures. The Combined Institutional Review Board at the University of Colorado at Anschutz approved the transfer and analysis of the de-identified samples collected in Sweden. The study protocol in Colorado (COMIRB number 20-1325) was determined Exempt/Non-Human Subject Research and was conducted in accordance with 45 CFR 46 and institutional policies. All data were pseudonymized to protect patient privacy.

### Taste evaluation

As mentioned, the human sense of taste is divided into five taste qualities: sweet, umami, bitter, sour and salty, (reviewed in:(Hellekant 2024; Lindemann 1996). To assess a subject’s ability to taste we used the quantitative and qualitative Waterless Empirical Taste Test (WETT^®^) by Sensonics (Haddon Heights, NJ, USA). The WETT^®^ test also measures the ability to taste each taste quality with an eight-point scale from zero (no taste) to a maximum of eight (all concentrations identified). The WETT^®^ measures the ability to identify various concentrations of the above five basic taste qualities with a minimum number of trials. It consists of 53 strips impregnated with one tastant (a substance that produces a taste sensation) representing a specific taste quality. A monomer cellulose pad (1 × 2.5 cm) is present on one side of each strip, containing either: dried <u>s</u>ucrose (sweet) (0.20, 0.10, 0.05, or 0.025 g/ml), citric acid (sour) (0.025, 0.05, 0.10, or 0.20 g/ml), sodium chloride (salty) (0.0313, 0.0625, 0.125, or 0.25 g/ml), caffeine (bitter) (0.011, 0.022, 0.044, or 0.088 g/ml), monosodium glutamate (umami; brothy) (0.017, 0.034, 0.068, or 0.135 g/ml), or blank control strips, with no added chemical stimulus. The subjects were instructed to place the appropriate strip on the tongue and move it along the surface of the tongue for at least 5–10 seconds. The subject indicated which taste sensation was perceived sött (sweet), bittert/beskt (bitter), surt (sour), buljongliknande smak/umami (umami), or salt (salty), or whether no taste was sensed by verbally reporting their response to be entered on the designated line of the response form. In the WETT^®^ test sequence, the concentrations of each stimulus are presented twice (eight total trials/tastant). In the first half of the test, the concentrations ascend from weak to strong with the different tastants presented randomly, with no tastant blanks included. In the second half of the test, the reverse presentation order is used, i.e., going from strong to weak concentrations. Scores range from 53 (all correct) to 13 (no tastes reported, corresponding to the number of control strips in the test). Each subject’s results were compared with normative values for each sex and age given in Doty et al (Doty *et al*. 2025).

### Biopsy procedures

Five to eight fungiform papillae (FP) were biopsied from each subject’s tongue and preserved for histological and molecular analyses. Prior to biopsy, the tongue was anesthetized with a Xylocaine spray (100 mg/ml Lidocaine with 10mg/spray dose), then blue or green food coloring was added to enhance the tongue’s fungiform papillae (FP). The tongue was held steady with a gauze, and biopsies of the FP were taken with a 3-mm sterile microscissor with a curved tip. The FP was cut one by one by holding down the scissor tip at the base of the chosen FP, which was removed with a single cut and put in fixative for histology or RNA*later*™ (Thermo Fisher Scientific Inc., Waltham, MA, USA). The biopsies for histological analysis were fixed in 4% buffered formaldehyde for at least four hours or overnight and then moved to phosphate-buffered saline (PBS) solution. The biopsies were stored at 4°C until shipping for analyses from Sweden to the United States for histopathological studies. The biopsies for RNA analysis were placed in RNA*later*™ stabilization solution and stored at room temperature overnight and then stored in the freezer at -18 °C initially and then at -80°C until analyses.

For all but two biopsy subjects, samples were taken immediately following the WETT test or within two months of the WETT test as indicated in Table 1. For the other two subjects (subjects SWE26 and 21), biopsies were taken four months following the WETT test. No biopsy was requested for subjects SWE1-6 or SWE10 because these subjects showed relatively normal total taste scores; subject SWE16 declined the biopsy procedure. For all other subjects (SWE7-9 ; SWE12-15, and SWE 18-29) biopsies were obtained without regard to taste score. Subject SWE17, who gave a biopsy sample, was a subject who recently tested positive for SARS-CoV-2 but reported no associated taste dysfunction. Histological and molecular data from this subject are included in our analyses, but the taste testing data are not included in the analysis of Long COVID taste testing which includes all subjects reporting taste dysfunction at the time the biopsy was taken and no biopsy cohorts.

### Histology

Upon receipt of fixed samples in Colorado, all specimens were cryoprotected in 20% Sucrose in 0.1M phosphate buffer overnight at 4°C. Then, the tissues were cryo-sectioned (12-14µm) onto Superfrost Plus microscope slides (Thermo Fisher Scientific, (Catalog# 1255015) and air-dried. Slides were rinsed in 0.1M PBS pH 7.2. Antigen retrieval was performed with 10 mM sodium citrate buffer of pH 6.0 at 85°C for 10 minutes before incubation in blocking solution (2% normal donkey serum, 1% bovine serum albumin, and 0.3% Triton in 0.1M PBS) for an hour at room temperature. Slides were then incubated for four days in specific primary antibodies diluted in blocking solution at 4°C including the following antisera:

The first two antisera were utilized in all cases, the latter two were used in some cases.

- Mouse anti-Tubulin β3 (clone TUJ1) at 1:1000 dilution (BioLegend, San Diego, CA, USA, catalog# 801202; RRID:AB_2313773) (for total innervation),
- Rabbit anti-PLCβ2 (PhosphoSolutions, Denver, CO, USA, catalog# 1640-PLC; RRID:AB_2801663), or guinea pig anti-PLCβ2 (PhosphoSolutions, AP8/24; RRID: AB_2910247) (for identification of Type II TRCs).
- Goat anti-GNAT3 at 1:500 dilution (Aviva Systems Biology, San Diego, CA, USA, catalog# OAEB00418; RRID:AB_10882823) (for TRCs expressing the taste-associated G-protein gustducin),
- Rabbit anti P2X3 (Alomone, Jersusalem, Israel, catalog #APR-016, RRID AB_2313760) (for labeling of taste bud-specific innervation).

After washing in 0.1M PBS, the slides were incubated two hours with fluorescent secondary antibodies, Alexa Fluor 488 donkey anti-mouse (Invitrogen, catalog# A21202; RRID:AB_141607), Alexa Fluor 568 donkey anti-rabbit (Invitrogen, catalog# A10042), Alexa Fluor 647 donkey anti-goat (Invitrogen, catalog# A21447; AB_2534017), Alexa Fluor 647 donkey anti-guinea pig (Jackson Immuno Research Laboratories, West Grove, PA, USA, catalog# 706605148; RRID: AB_2340476) or Donkey anti-Rabbit IgG (H+L) Highly Cross-Adsorbed Secondary Antibody, Alexa Fluor™ 647 (ThermoFisher Catalog # A-31573; AB_2536183) at 1:800 dilution each at room temperature. Lastly, the slides were washed twice for 10 minutes each in 0.1M PBS and then once for 10 minutes in 0.05M PBS before coverslip slides with DAPI Fluoromount-G (SouthernBiotech, Birmingham, AL, USA, catalog# 010020).

### RNA extraction

The papilla tissue samples for PCR analysis were stored in RNA*later*™ at -20°C. RNA was extracted with the RNeasy fibrous tissue mini kit (Qiagen, Hilden, Germany). A small piece of the biopsy was taken with a pipet tip into a 1.5 ml tube containing 300 µl of RLT buffer and mashed manually with a micro pestle. The pestle was rinsed when pipetting 590 µl of RNase free water into the tube. The kit protocol was followed, and a DNase treatment was performed with a DNA-free DNA removal kit (Invitrogen) according to their protocol. The resulting RNA concentrations and quality were checked with Nanodrop 8000 (Thermo Fisher Scientific) and TapeStation High Sensitivity RNA tape (Agilent, Santa Clara, CA, USA).

### Reverse transcription

High-Capacity cDNA Reverse Transcriptase Kit (Applied Biosystems, Waltham, MA, USA) was used to obtain cDNA from the RNA samples. 100 ng of RNA per sample, according to TapeStation concentrations, was taken into the reverse transcription reaction. The kit protocol was followed. The reverse transcription was performed in duplicates to obtain enough product for all the qPCR runs.

### qPCR

Quantitative PCR (qPCR) was performed with TaqMan (Applied Biosystems) assays for *PLC*β*2* (Hs01080542), *NCAM1* (Hs00941821), and *TAS1R3* (Hs01026531) as well as for two reference genes *RPLP0* (Hs99999902) and *GAPDH* (Hs02758991). TaqMan Gene Expression Master Mix (Applied Biosystems) was used according to its protocol, and 10 ng of cDNA was added as template. All samples were run in triplicate per gene, including a negative control. One gene per plate was run in the QuantStudio 5 (Applied Biosystems). Average Ct and standard deviation were calculated for each gene and then normalized (Delta CT =Ct(gene of interest) – Ct(reference gene)) against the reference gene *GAPDH*. The reference gene *GAPDH* gave more consistent normalization than *RPLP0*, and was therefore selected as a reference for the genes of interest.

### SARS-CoV-2 test

A PCR test for the presence of SARS-CoV-2 virus in the tissue samples was performed in triplicate, including two assays that target SARS-CoV-2 as well as negative control samples. TaqMan microbe detection assay (Vi07918637) for SARS-CoV2N and TaqPath Blastopore Microbial Detection Master Mix (Applied Biosystems) were used according to their protocol. Positive control was TruMark Respiratory Panel 2.0 Amplification control (Applied Biosystems). The qPCR was run in the QuantStudio 5 (Applied Biosystems). None of the biopsied tissue samples tested showed the presence of SARS-CoV-2 virus, although one subject (#17) had an active SARS-CoV-2infection as determined by a positive nasal swab test. Absence of detectable SARS-CoV-2 virus in oral or salivary samples despite positive detection of virus in nasal swabs is not uncommon and is described in other studies (Fernandes *et al*. 2021; Goodall *et al*. 2022).

### Statistical analysis

Parametric statistical calculations were carried out by Excel (Microsoft) or with calculators given at www.socscistatistics.com. Both calculators gave identical values. (Ho *et al*. 2019) For comparisons (Ho *et al*. 2019)across samples, we utilized Pearson’s Correlation co-efficient or Fisher’s Exact test for small sample sizes. For comparison of test percentile means against the expected 50% percentile distribution, we employed a one-sample t-test. A criterion value of p < 0.05 was set for statistical significance in all calculations.

Means, standard deviations, and ranges were used to describe continuous variables such as age, time from infection to testing, and taste scores. Categorical variables like sex and symptom presence were summarized using counts and percentages.

Because taste ability varies substantially by sex and age, raw taste scores were normalized by sex and age using reference data from Doty et al. (2025), which included 1,392 healthy individuals. Scores for individual tastes below the 10% compared to sex and age matched populations in Doty et al (2025) were considered potentially hypoguesic per the criterion set in (Landis *et al*. 2009) and are highlighted in red in Table 1.

The binomial test or Chi-square tests (χ²) were used to compare (categorical data) the frequency of abnormal taste scores across groups. The results for Chi-square tests are reported as χ² (degree of freedom (df), sample size) = chi-square value, p-value. Pearson’s Correlation (r) was used to assess linear relationships between molecular markers (qPCR results), histological features, and taste scores. The results were reported as (df) sample size=correlation co-efficient, p-value. (Schober *et al*. 2018).

## RESULTS

### Study population

All 28 subjects (24 women and 4 men) reported normal taste before the first confirmed SARS-CoV-2 infection but, except for subject SWE17, altered or diminished taste thereafter . The mean age at testing was 49.9 years (SD+/-1.2) and ranged between 30 and 71 years. Most subjects contracted COVID-19 in early 2020. Two subjects (SWE19 & 28) experienced a second infection in 2022, shortly before undergoing taste testing and biopsy. Considering only the first infection, the time from initial SARS-CoV-2 infection to taste testing ranged from 13.5 months to 27.0 months (mean of 21.0 months).

### Psychophysics

Because taste ability varies substantially between individuals according to sex, age, and health status, we normalized all raw taste scores and overall WETT score against values given for 1,392 normal subjects (Doty *et al*. 2025). Although all subjects, except one (subject #SWE17), reported compromised taste abilities in clinical reporting, only 3 of 28 tested subjects scored below the 10th percentile in overall taste ability consistent with overall scores in a population of normal subjects (Doty *et al*. 2021) (see Table 1) and consistent with previous reports of no demonstrable long-term taste loss in broad post-covid populations following SARS-CoV-2 infection (Boscolo-Rizzo *et al*. 2023a; Sharetts *et al*. 2024). Nonetheless, 14 of the 27 of our Long Covid dysguesia subjects showed significant deficiency (10th percentile or lower) in one or more individual taste qualities (values in red shown in Table 1) which is significantly more than expected in a normal population (binomial test 24 of a total of 135 individual taste qualities; p = 0. 002). sex(Doty *et al*. 2021) One subject, who was also anosmic, (SWE29) gave no correct responses to any tastant, even at the highest concentration suggesting complete aguesia. Even subjects with a near-average total WETT score (percentiled by sex and age group) were unable to detect specific tastes, e.g. subjects SWE8 (F, age 64; WETT Score 32; 45%) and SWE9 (M, age 49; WETT Score 30; 54%) were unable to detect bitter or umami. Although individuals with a profound overall deficit also showed specific loss of one or more taste qualities (e.g. subject SWE7 [16%] showed no response to sweet, bitter or umami, but good response to sour and salty), the pattern of loss was not duplicated by other subjects (e.g. subject SWE24 [7%] with good responses to sweet, sour and salty, but total loss for bitter and umami).

The distribution of loss between subjects was uneven with sour standing out as the least affected quality (only one score below the 10%). This may reflect the detection of “sour” being a multisensory experience involving chemesthetic as well as gustatory modalities (Ohkuri *et al*. 2012; Yu *et al*. 2020; Zhang *et al*. 2019). In contrast, in acute COVID-19, sour taste is reported to be most impacted during the acute phase of the illness (Karmous *et al*. 2023; Singer-Cornelius *et al*. 2021).

More importantly, as a group, the PLCβ2-mediated taste qualities (sweet, umami, bitter) showed total loss (score of 0) as well as scores below 10% significantly more than the other taste qualities (sour, salty) (*X*^2^(1, *N* = 135) = 4.47, *p* = .03). These findings suggest that Long Covid taste disturbance may preferentially target the PLCß2-mediated taste qualities transduced by Type 2 taste receptor cells.

### Quantitative PCR Findings

RNA prepared from taste papillae from all biopsied subjects, including SWE17, was analyzed for expression levels of three taste-related mRNAs encoding: 1) phospholipase C beta 2 (PLCß2) a marker of Type II cells, 2)Taste receptor member *TAS1R3*, coding for a receptor component for both sweet and umami tastes and expressed in a subset of Type II taste cells; and 3) neural adhesion molecule 1 (*NCAM1)*, a marker of total innervation as well as Type III TCs. All data were normalized against glyceraldehyde-3-phosphate hydrogenase (*GAPDH)* mRNA, yielding Delta CT values which are expressed as a fold-change compared to the values from subject SWE17 who described no taste dysfunction. These relative expression values are given in Table 2 along with Taste Scores and computed average taste score for sweet and umami (Ave Sweet-Umami) – the Tas1R1-related taste qualities -- for comparison.

**Table 2:** Relative expression of taste-related mRNAs from qPCR of fungiform papillae. Subject SWE 17, who reported no taste disturbances, served as our comparator. Taste test values as shown in Table 1 are included for comparison. For each subject, the average of the scores for sweet and umami taste, tastes reliant on Tas1R3, are shown as well.

| Subject # | taste test value | relative expression |  |  | Ave PLC + Tas1R3 | ave sweet/umami |
| --- | --- | --- | --- | --- | --- | --- |
|  |  | PLCb2 | TAS1R3 | NCAM1 |  |  |
| **SWE 17** | **53** | **1.00** | **1.00** | **1.00** | **1.00** | **8** |
| SWE 7 | 24 | 0.15 | 0.11 | 0.96 | 0.13 | 0 |
| SWE 8 | 32 | 0.66 | 0.34 | 0.60 | 0.47 | 3 |
| SWE 9 | 30 | 0.29 | 0.42 | 0.40 | 0.35 | 0.5 |
| SWE 12 | 35 | 0.78 | 0.51 | 1.17 | 0.63 | 3 |
| SWE 13 | 22 | 0.30 | 0.78 | 1.01 | 0.49 | 0.5 |
| SWE 14 | 40 | 0.53 | 1.60 | 1.21 | 0.92 | 5.5 |
| SWE 15 | 34 | 0.56 | 0.77 | 0.67 | 0.66 | 5 |
| SWE 18 | 32 | 0.14 | 0.30 | 0.15 | 0.20 | 4 |
| SWE 19 | 36 | 0.16 | 0.76 | 0.08 | 0.35 | 5.5 |
| SWE 20 | 45 | 1.64 | 0.80 | 0.86 | 1.14 | 4.5 |
| SWE 21 | 34 | 0.27 | 0.23 | 0.34 | 0.25 | 2.5 |
| SWE 22 | 26 | 0.36 | 0.28 | 0.52 | 0.32 | 2.5 |
| SWE 23 | 39 | 0.64 | 1.14 | 1.96 | 0.86 | 4 |
| SWE 24 | 17 | 0.62 | 1.16 | 1.33 | 0.84 | 2.5 |
| SWE 25 | 19 | 0.19 | 0.17 | 0.52 | 0.18 | 3 |
| SWE 26 | 37 | 0.48 | 0.59 | 0.57 | 0.54 | 3 |
| SWE 27 | 25 | 1.08 | 0.77 | 1.64 | 0.91 | 0 |
| SWE 28 | 36 | 0.44 | 0.54 | 0.32 | 0.49 | 4.5 |
| SWE 29 | 13 | 0.22 | 0.15 | 0.59 | 0.18 | 0 |

**Table 3:**
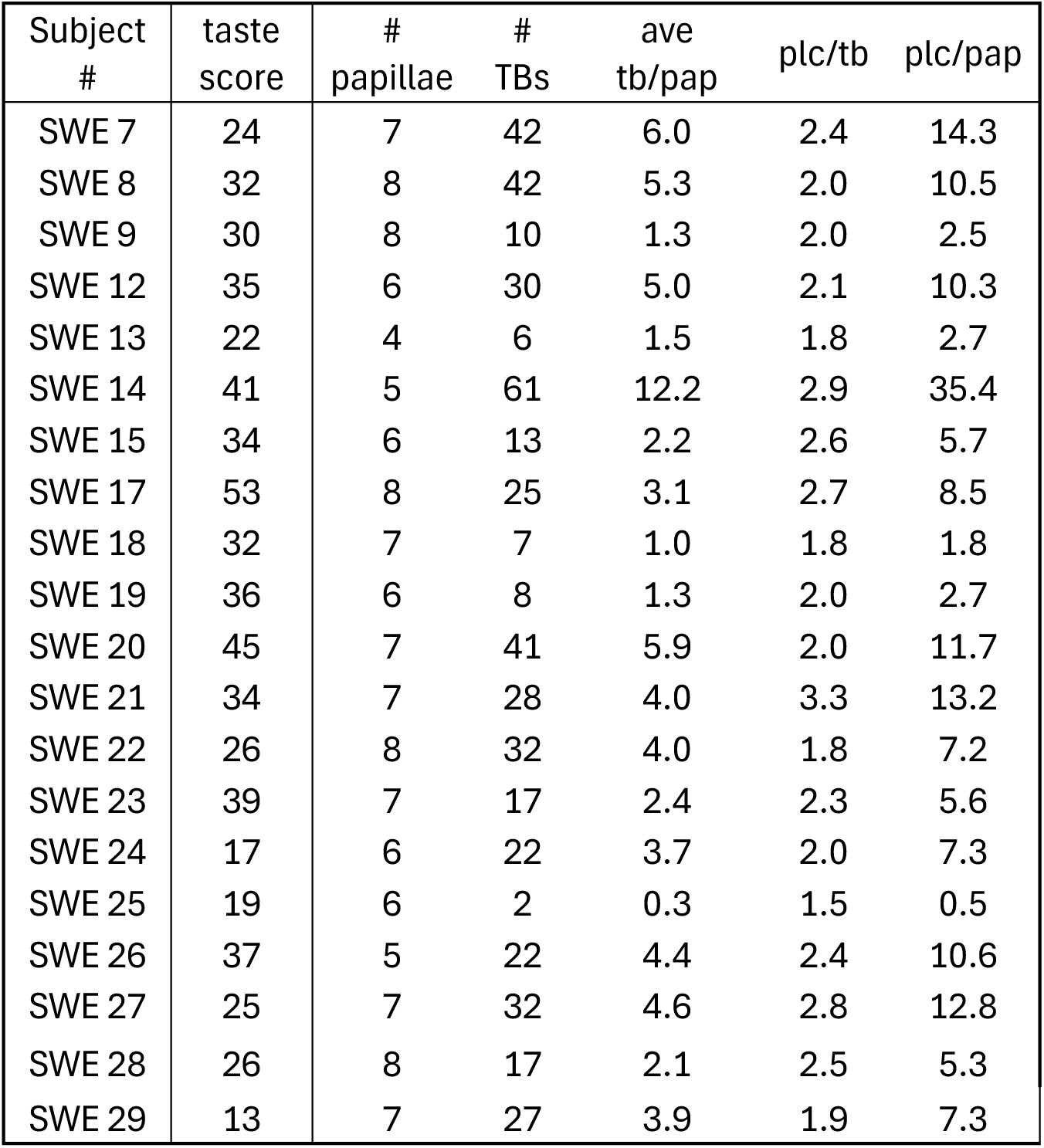
Counts of papillae sampled, number of taste buds and average *PLCß2*+ cells per taste bud and papilla in each case. No significant correlation was found between taste score and average number of taste buds per papilla nor of density of *PLCß2*+ cells per taste bud or papilla. The histological counts and density of *PLCß2*+ cells also did not correlate with any PCR values. # TBs, number of taste buds; ave tb/pap, average number of TBs per papilla; PLC/tb, average number of PLC+ cells per taste bud; PLC/pap, average number of PLC cells per papilla.

The expression levels for *PLCß2* and *TAS1R3* mRNAs, both encoding components of the taste transduction cascade in Type II TCs, were highly correlated (r(18) = 0.64, p=0.002) as expected based on molecular features of Type II taste cells in mice. Expression of *NCAM* mRNA was not significantly correlated with either *PLCß2* or *Tas1R3* mRNAs.

Comparing the qPCR data from all biopsy subjects to the taste scores from the same subjects shows a significant correlation of overall taste score to *PLCß2* (r(18) = -0.51, p= 0.023) and substantial correlation with *TAS1R3* (r(18) =--0.43, p= 0.056; although not significant at p<0.05 level). This relationship is shown graphically in Fig. 1. Conversely, expression levels of *NCAM* mRNA showed no substantial correlation with taste test values (r(18) = 0.035, p>0.05).

**Figure 1:**
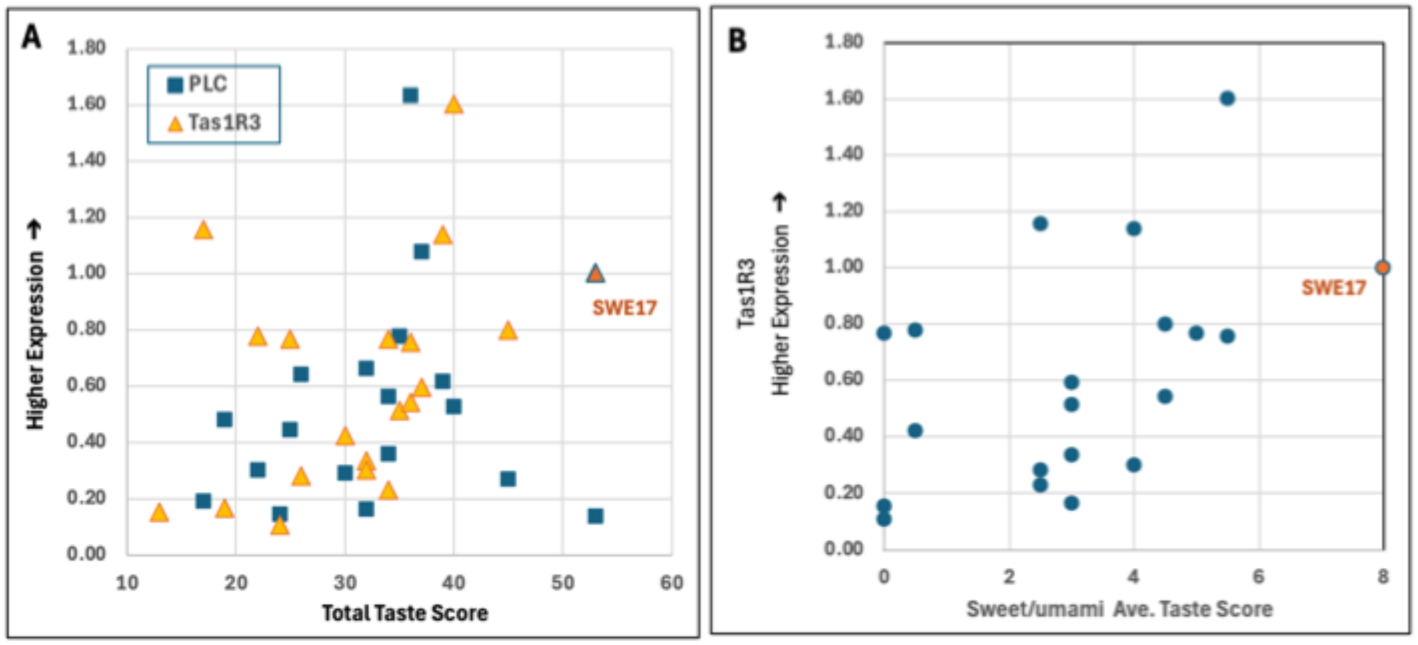
Correlation of relative *mRNA* expression levels with taste scores. The point indicated in orange is the data from Subject SWE17 who was used for comparison. A) The mRNA levels for *PLCß2* (r(18)=-.51, p= .023) and *TAS1R3*(r(18)=.43, p= .056) correlated with the overall taste score for each subject. B) The mRNA level for *TAS1R3* correlates with the combined taste score for the related taste qualities (Sweet and brothy) (r(18)=.51, p= .02). Alt Text: Graphs showing the correlation between Taste Score and PCR analysis of taste-related markers.

Since both sweet and umami require the TAS1R3 receptor subunit as well as PLCß2, we compared the expression level of these mRNAs to the average of taste scores for sweet and umami for each subject. A significant correlation exists between the average of the related taste scores for sweet and umami with theDelta CT relative expression values for *TAS1R3* (r(18)= 0.45, p=0.046) and *PLCß2* (r(18)= 0.55, p=0.01). These expression levels were, however, not significantly correlated with expression levels of the individual tastes: sweet or umami.

### Histological Findings

Fungiform samples from the subjects varied between 4 and 8 papillae, with a median of 7. Approximately half of the papillae had no identifiable TBs, as reported for human fungiform papillae in previous literature e.g. (Miller 1988). Normal-appearing taste buds were identified in all subjects, including subject 29, who reported no taste sensibility. While relatively normal TBs occur in samples of all subjects (See Fig. 2), many subjects displayed disorganized TBs or unusual isolated PLCß2+ cells which were never observed in TBs from normal subjects (High and Finger 2025)

**Figure 2:**
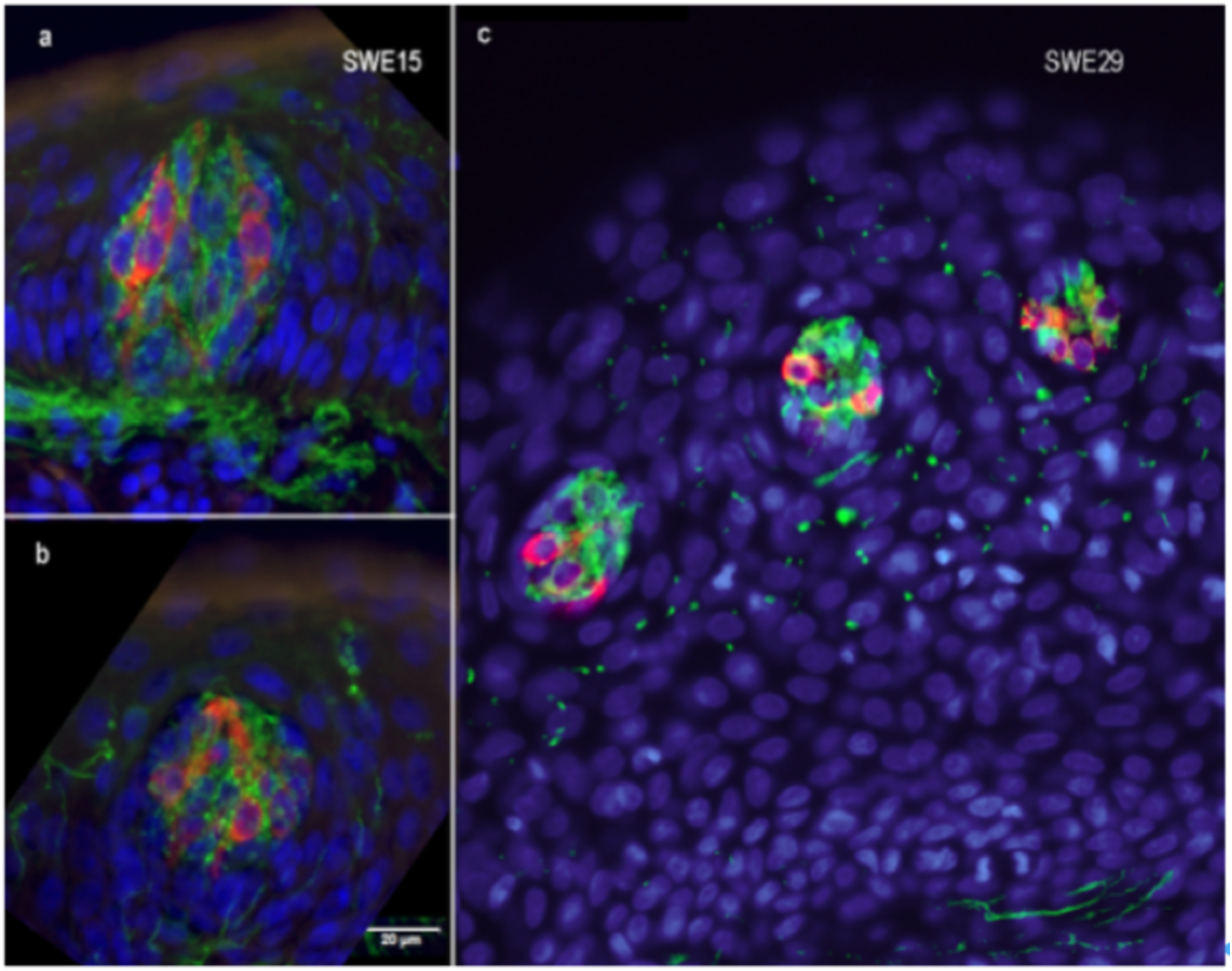
Appearance of normal taste buds sectioned in different planes from different subjects. Red = PLCß2 immunoreactivity showing Type II taste cells; Green = TUJ1 immunoreactivity for nerve fibers. Typical taste buds have more than one PLCß2+ cell surrounded by a dense network of TUJ1- stained nerve fibers. Green profiles outside of taste buds are extragemmal nerve fibers typical of lingual epithelium (Moayedi *et al*. 2021). **a, b.** Taste buds sectioned longitudinally appear similar in shape to garlic bulbs or onions (from Subject (SWE) 15. **c)** Taste buds sectioned perpendicular to their long axis appear circular (from subject (SWE) 29 who reported a total absence of taste perception (taste score of 13, showing no correct responses to any tastant). Scale bar in panel (b) applies to all panels. Alt Text: Images of 3 micrographs showing the normal appearance of taste buds in our samples. Multiple PLCs2-positive taste cells within each bud are embraced by a dense nerve plexus.

The average number of TB profiles per papilla was 3.6 (+/-2.6 s.d.). Because of the significant variance in the number of TB profiles observed in a single section through a papilla, we calculated the average number of TB profiles per papilla from all papillae to measure the taste bud population for each subject. In addition, we counted the number of PLCß2+ taste cells within each section to calculate the average proportion of PLCß2+ cells per TB. The taste score for each subject was not correlated significantly with the average number of TB profiles per papillae (r(18) = 0.268, p = 0.25), with the average number of PLCß2+ cells per papilla (r(18) - 0.35, p= 0.15), nor with the average number of PLCß2+ cells per TB (r(18) = 0.427, p = .06). As shown in Fig. 3, abnormal TBs or irregular PLCß2+ cells were especially noteworthy in 7 of our 19 post-covid biopsy subjects (SWE12 (Taste Score 54%), 13(21%), 22 (21%), 24 (8 %), 25 (4%), 28 (44%) and 29 (1%)). This incidence is significantly different (Fisher’s exact test p <0.03) than in tissues obtained from normal subjects where dispersed PLCß2+ cells were never observed in fungiform taste buds from 12 subjects.

**Figure 3:**
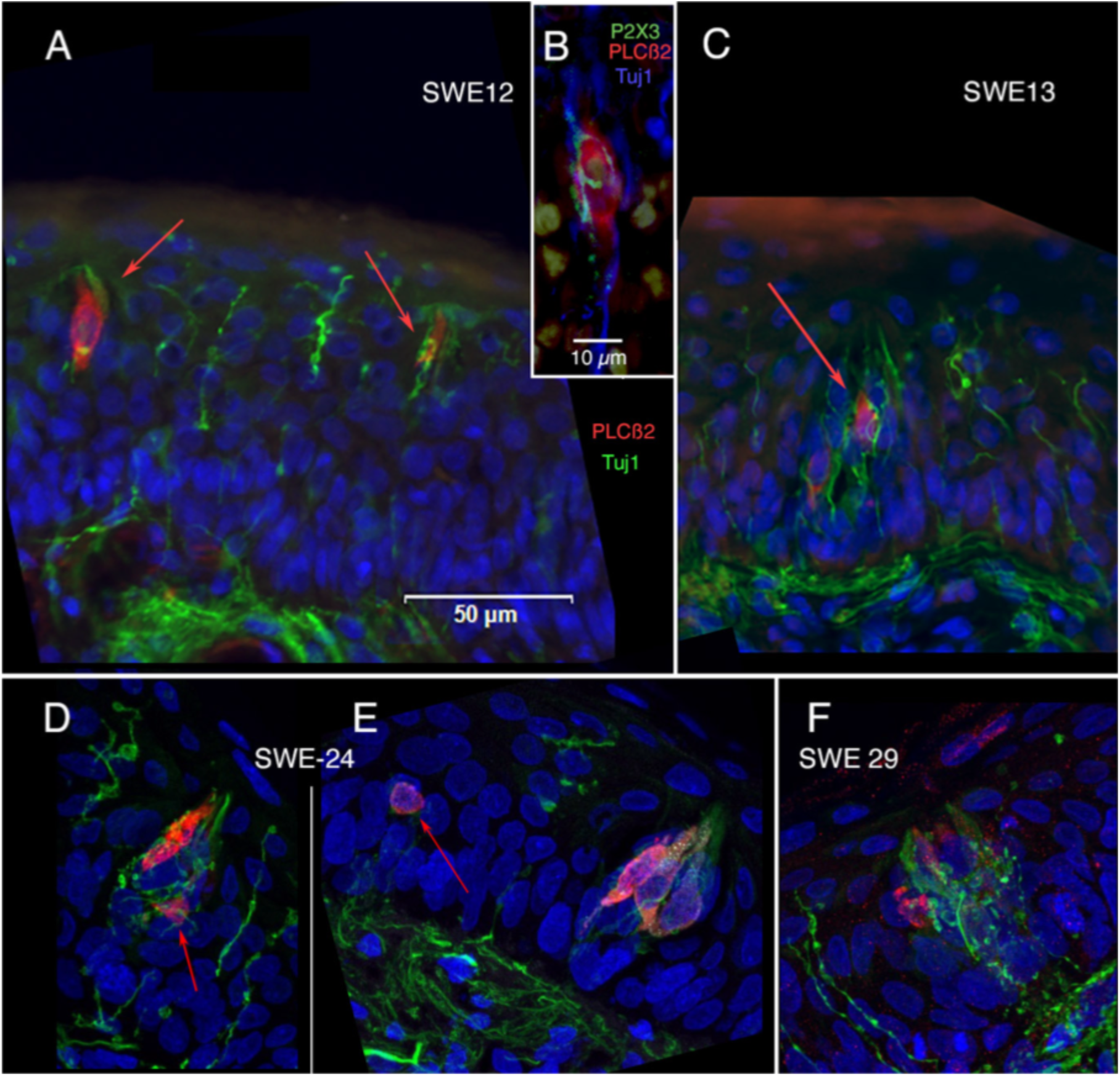
Examples of abnormal PLCß2+ taste-like cells from 4 cases. (A & B). Subject (SWE) 12 had a taste score of 35 (54%). (C) From subject (SWE) 13 who had a score of 22 (21%). In both cases (A & C), innervated PLCß2+ (red arrows) cells occur in relative isolation within the epithelium. Innervation of these cells (green: TUJ1) is sparse unlike innervation of taste buds (cf Fig. 2). These cells may represent very small taste buds or simply PLCß2+ cells lying outside of a taste bud structure. Panel B shows a higher magnification of one such isolated PLCß2+ cell and its innervation stained for the presence of P2X3, a neural receptor predominantly on nerve fibers that innervate taste buds. Panels D-E are from case (SWE) 24 with a taste score of 17 (4%) showing examples of aberrant PLCß2+ cells (red arrows) outside of a taste bud structure. Panel F: A disorganized taste bud from case SWE 29 who reported a total absence of taste perception (taste score of 13, showing no correct responses to any tastant). The heavy innervation is typical of taste buds, but the cellular structure revealed by PLC-staining (red) does not show typical taste bud organization. Alt Text: Images showing abnormal isolated PLC-positive cells and abnormal taste buds within the epithelium of our subjects.

## DISCUSSION

### Morbidity Changes in COVID-19 caused by SARS-CoV-2 Variants

Previous studies found that the frequency of chemosensory symptoms, including taste loss, varies across COVID-19 illness caused by infection of different SARS-CoV-2 variants. Early variants, such as Wild type and Alpha, were more commonly associated with taste and smell loss, while later emerging variants, such as Omicron, caused lower levels of taste loss (Reiter *et al*. 2023). Most subjects included in our study were primarily infected during the first wave of the COVID-19 pandemic, and their reported taste disturbances fit well with this pattern of greater taste loss early in the pandemic.

Published epidemiological data further illustrate this trend. Among individuals infected with the Wild-type strain, 241 out of 1,141 cases (21%) reported smell and/or taste disorders. For the Alpha variant, 223 out of 1265 cases (18%), for later variants such as Delta, 480 out of 1516 cases (32%), and for Omicron, only 317 out of 7431 cases (4%) presented with smell and/or taste disorders (Ota *et al*. 2025). This supports the hypothesis that the variant-dependent tropism and immune response may influence the degree and duration of taste dysfunction.

### On the role of the ACE2 receptor

Relatively early in the COVID-19 pandemic, it was suggested that SARS-CoV-2 infects oral tissue (Huang *et al*. 2021b). Doyle et al. (Doyle *et al*. 2021) reported that ACE2 was expressed by Type II TRCs, and further demonstrated the presence of SARS-CoV-2 in Type II TRCs, which mediate sweet, umami and bitter sensations, but not within the other taste cell types. This likely explains why subjects in that study did not taste the “sweetness of chocolate and bitter coffee” and instead reported that it “tasted like mud” .

Research during the COVID-19 pandemic has highlighted the potential impact of SARS-CoV-2 on TBs, particularly through its interaction with ACE2 receptors (Huang *et al*. 2021a; Yao *et al*. 2023). In patients experiencing taste disturbances following COVID-19 infection, SARS-CoV-2 was detected in basal and suprabasal layers of fungiform papillae for extended periods. In one subject, virus was detected up to 63 weeks after the initial diagnosis, coincident with disrupted nerve fibers and altered taste bud cytoarchitecture (Yao *et al*. 2023).

Recovery of taste, and re-establishment of normal epithelial cytoarchitecture, coincided with the disappearance of the virus from the taste epithelium (Yao *et al*. 2023). These findings suggest that prolonged viral presence or the possibility of reinfection and the associated immune response may impair the renewal and functionality of taste buds in Long COVID. In our subjects, a sensitive commercial qPCR test showed that none of the tissue samples were positive for the SARS-CoV-2 viral RNA suggesting that taste dysfunction may persist even after the virus is no longer detectable in the peripheral taste system. This offers the potential that persistent taste dysfunction in Long Covid may outlast the presence of the virus itself.

### Why loss of sweet-bitter-umami tasting ability is more frequent than the loss of other taste qualities

A previous WETT study involving 340 long-term (1-year) COVID-19 subjects and 434 non-COVID controls (Sharetts *et al*. 2024) found no statistically significant differences in overall taste function between groups. Similarly, a psychophysical taste-strip study (Boscolo-Rizzo *et al*. 2023a) of 62 post-Covid subjects who reported an altered sense of taste or smell one year after infection, found no statistically significant evidence for ongoing taste loss by two years after infection, although subjects still scored modestly lower than the normal population. Similarly, our subjects show non-significant, slightly lower than average total taste scores (46.5%±25.1 s.d.) compared to a normal population. It should be noted that our study specifically targeted subjects reporting ongoing taste loss, not general chemosensory disruption, two years after COVID-19 infection, preventing direct comparison of our results with those surveyed from the broader population surveyed in Sharetts et al. (2024) or when patients with shorter-term chemosensory loss were tested (Boscolo-Rizzo *et al*. 2023b).

Despite the lack of measurable overall taste loss in our subjects, our psychophysical results do indicate a significant disruption of taste with a tendency toward preferential impact on sweet, bitter, and umami -- the PLCβ2-related taste qualities (Zhang *et al*. 2003). These taste modalities all are detected by the Type II taste cells which are targeted by SARS-CoV-2 virus (Doyle *et al*. 2021).

Taste dysfunction may result from disruption of taste-related intracellular transduction cascades or damage to taste buds (TBs), their receptor cells (TRCs) or the gustatory nerves. (Liman 2020; Liman and Kinnamon 2021; Zhang et al. 2003; Zhao et al. 2003). The fact that the SARS-CoV-2 can directly infect Type II TRCs may explain why deficits in sweet, umami and bitter are more frequent than for sour and salt. In our study, however, viral RNA was not detectable in taste tissues and there was no evidence of wholesale destruction or abnormality in Type II, PLCβ2+, taste cells. Nonetheless, in our subjects overall tasting ability correlates positively with the amount of PLCβ2 and TAS1R3 mRNAs present in the tissue. Curiously, a previous study (Donadoni et al. 2023) of the consequence of SARS-CoV-2 infection of taste cells in an in vitro culture system showed increases in PLCβ2 expression with infection. It is unclear how the acute in vitro model may relate to the long-term post-covid conditions reflected in our study. Hypothetically, the long-lasting effect could be the result of an epigenetic mechanism called inflammatory memory (Naik et al. 2017). This is described in several inflammatory diseases and the effect is manifested as a sustained modification of the chromatin structure. This leaves open the question how prior viral infection might induce long-lasting deficits in taste.

In the olfactory epithelium, the virus infects supporting cells, but not the olfactory receptor cells (Chen *et al*. 2020). Nonetheless, the olfactory receptor cells responded to the viral infection of the tissue by downregulating expression of olfactory receptors and downstream signaling molecules while upregulating expression of viral response proteins (Chen *et al*. 2019; Zazhytska *et al*. 2022). The transduction cascade is re-established in the receptor cells once the viral infection has been cleared (Zazhytska *et al*. 2022). While a similar phenomenon may occur in taste buds during viral infection, it is unlikely such a reaction would persist in the tissue after the virus is cleared and all the taste cells have been replaced during the course of normal taste cell turnover which requires a few months at most (Perea-Martinez *et al*. 2013). At the time of our testing, up to two years after initial infection, none of the taste cells present at that time would have been present at the time of viral infection.

Furthermore, we do note that some subjects are unable to detect non PLCβ2-dependent taste qualities, i.e. sour and salty. This finding may reflect inabilities typical of a normal population but might suggest that non-PLCβ2 related mechanisms may also be at work underlying taste dysfunction in Long COVID. One possibility is that the infection alters the function in taste nerves. Indeed, evidence exists for widespread neural dysfunction in Long COVID (Daia *et al*. 2021; McFarland *et al*. 2021; O’Neill *et al*. 2023; Otte *et al*. 2022; Rogn *et al*. 2024; Stepien and Pastuszak 2023). If so, then some of the observed taste loss might be attributable to loss of transmission in the taste nerves rather than disruption of the taste epithelium. This could explain the total loss of taste perception in subject SWE29 despite the presence of normal-appearing taste buds.

### Other factors that may have influenced taste during the disease process of COVID 19

Recent studies (Finlay *et al*. 2022; Peluso *et al*. 2024a; Peluso *et al*. 2023) broaden the perspective by suggesting that immune dysregulation and T cell dysfunction may play significant roles in the long-lasting effects of COVID-19. Whole-body positron emission tomography imaging with [^18^F]F-AraG, a tracer that selectively tags activated T cells, showed that subjects with Long COVID had specific tissues enriched for activated T cells compared to individuals without this condition (Peluso *et al*. 2024b). All five subjects in that study showed the presence of intracellular SARS-CoV-2 spike protein-encoding RNA in lamina propria tissue up to 676 days after initial infection with SARS-CoV-2. This offers evidence of viral persistance or of multiple independent SARS-CoV-2 infections which could underlie long-term immunologic perturbations for up to two years after an initial infection. Whether a similar prolonged immunological response occurs in taste tissue is unknown although Yao et al (Yao *et al*. 2023) report long term (>6 months) inflammatory response when virus remains in the tissue. We did not, however, detect any virus by PCR in our tissue samples from patients more than one year post infection. Other investigators reported an association of elevated serum concentrations of IL-10 and IL-1β with persistent taste dysfunction for bitter and sweet qualities, respectively, following Covid-19 (Karmous *et al*. 2023; Locatello *et al*. 2021).

### On the relation between histological changes of TB by COVID-19 and taste changes

A particularly intriguing aspect of our study is that all subjects, including those with severe deficits, displayed structurally intact taste buds despite the loss of taste function. However, some dystrophic taste buds and isolated PLCß2-expressing cells were observed in many subjects. In mice, disruption of molecular signals between nerves and taste buds can result in formation of aberrant PLCß2-expressing cells outside of the taste buds themselves (Castillo *et al*. 2014). This finding raises important questions about the disconnect between histological integrity and functional impairment. One possible explanation is that taste bud dystrophy, rather than outright loss of taste buds, may be sufficient to alter taste sensitivity by impairing cellular turnover or disrupting synaptic communication with gustatory neurons (Wang *et al*. 2023). While intact taste buds might suggest some functional resilience, the significant decrease in PLCß2-dependent signaling suggests that molecular-level dysregulation may underlie the persistent taste dysfunction. In addition, the discrepancy between PLCß2 expression and taste bud morphology, especially in Subject 29 who shows complete taste loss, suggests dysfunction in peripheral nerves or at higher levels of the neuraxis.

### Clinical relevance

Understanding the mechanisms behind taste loss is clinically relevant, as chemosensory loss negatively impacts nutritional intake, emotional well-being, and quality of life. For survival, taste ability is more important than both hearing and vision. A multidisciplinary mindset is crucial for appropriate diagnostics and for providing patients with appropriate treatment and underscore the importance of developing targeted therapies to alleviate the burden of long-term taste dysfunction in COVID-19 survivors.

Here we present an integrative approach combining psychophysical taste testing, RNA expression analyses using markers for TRC type II and III, and histological evaluation of taste tissue in subjects with Long COVID. This multidimensional analysis provides a more comprehensive understanding of Long COVID taste dysfunction. While this study focuses on the loss of Type II TRC, PLCß2-dependent taste qualities (sweet, umami, bitter), the loss of other taste qualities in a limited number of subjects suggests that multiple mechanisms may underlie the sensory loss.

### Limitations & Conclusions

The study reported here is a retrospective clinical study of post-Covid patients and like other case series studies has no control group although we offer a comparison to a pre-pandemic dataset on gustatory phenotypes in healthy individuals (Doty *et al*. 2025). Another limitation is that our study includes only a single time point measurement of taste function in a limited number of subjects. This limits the statistical power of the study, but nonetheless we do find highly significant correlated results even in our modest number of subjects. Furthermore, the study relies on self-reported accounts of taste loss coinciding with COVID-19 onset, but baseline taste function data before infection was not possible perhaps reducing the generalizability of the findings.

Despite these apparent limitations, our study presents important new quantitative psychophysical data showing significant disruption in detection of specific taste qualities in a subset of subjects reporting long-term post Covid taste dysfunction. The taste qualities most impacted are those detected by Type II taste cells which utilize a PLCβ2 dependent signaling cascade for transduction of sweet, umami and bitter qualities. Importantly, our study also shows a correlated decrease in mRNA levels for two elements of the Type II cell transduction cascade for these qualities –- the first evidence for correlative changes in molecular parameters tracking psychophysical taste functions. Finally, despite the well-documented recovery of post-Covid taste function in most patients, our study offers quantitative evidence for prolonged disruption of taste abilities in a small subset of post-Covid subjects.

## ACKNOWLEDGEMENTS

This work was supported by grants 3R01DC014728-05S1 to TEF awarded by the National Institute on Deafness and Other Communication Disorders of the National Institutes of Health, by Åke Wiberg Stiftelsen grant 2023 to M.K. Governmental funding of clinical research within the Uppsala university Hospital (ALF), and by internal funding of the Department of Animal Biosciences, Swedish University of Agricultural Sciences.

## AUTHOR CONTRIBUTIONS

G. H. and T.E.F. conceived this research and along with G.L.and G.A. designed the research plan. G.H. & G.L. wrote the ethical proposal to Etikprövningsmyndigheten. H.M. provided subjects, performed the WETT tests and took the biopsies. M.A.K. provided subjects and with A.A. assisted in analysis of the subject population. T.E.F. and M.L. designed and performed the histopathology and data analysis. G.A. and T.V. designed and analyzed the quantitative PCR experiments. T.V. performed RNA extraction and quantitative PCR experiments. T.E.F. performed statistical analysis of the quantitative data and compared results to those in the Doty et al (2015) report. G.L., G.H., G.A., H.M., and T.E.F. provided funding for the project. G.H., T.E.F., G.A., and M.A.K. prepared the manuscript and all authors have discussed and revised the manuscript.

### Use of AI

This study utilized Microsoft Copilot (GPT-4) accessed through a secure university-managed Microsoft Teams environment. The tool was utilized in the process of editing and checking calculations but was not used to generate any original content. All outputs were reviewed by the lead author.

## Data Availability Statement

The data underlying this article will be shared on reasonable request to the corresponding authors.

---

**Supplementary Figure 1:**
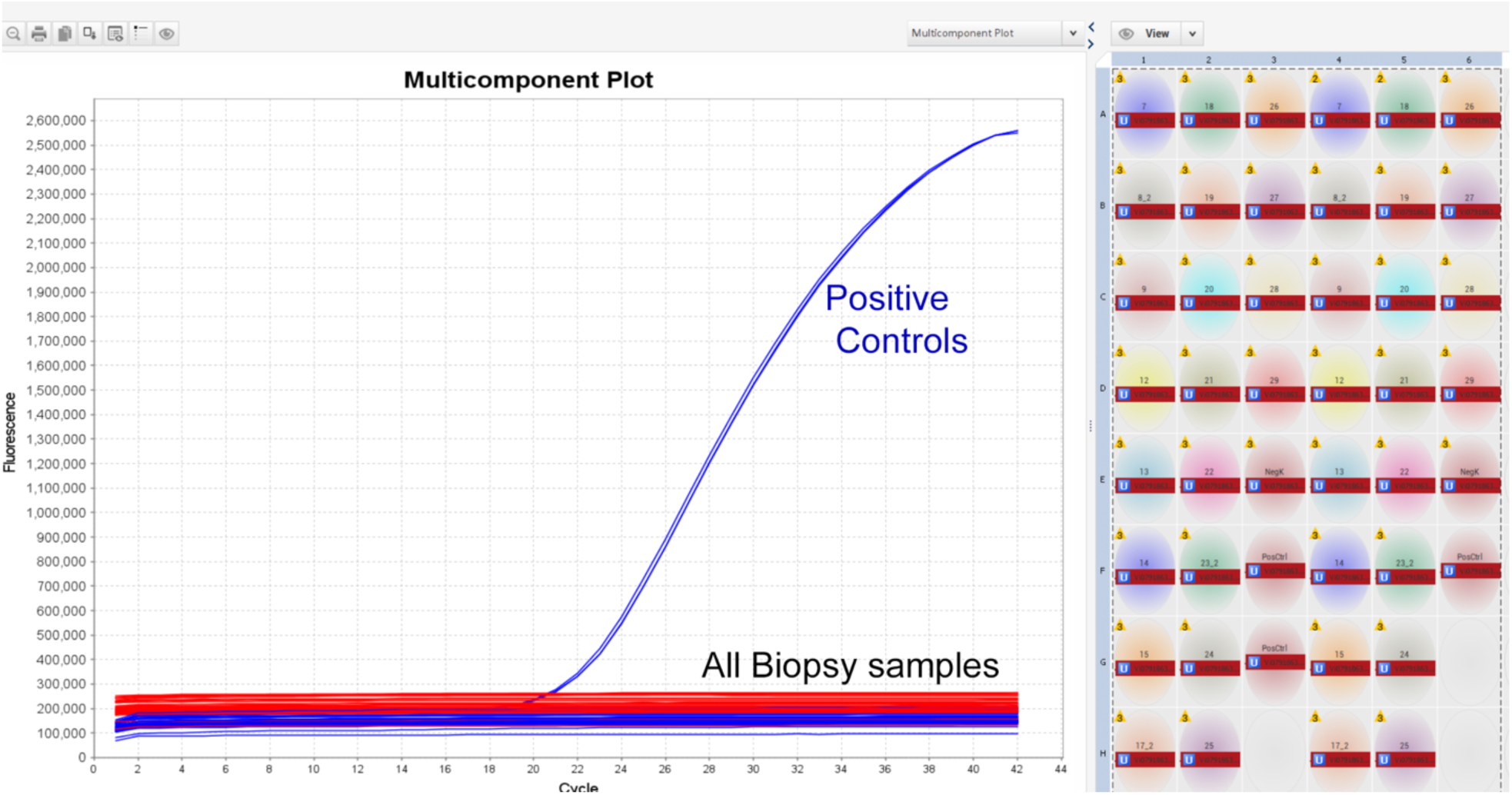
qPCR for Covid mRNA showing positive controls. No samples from any subjects show evidence of reaction for Sars-CoV2 virus.

**Supplementary Table 1:** WETT taste test values shown as a percentile values as normalized to the data in Doty et al 2025.

| Subject Number |  | Sweet | PLC Tastes<br>Umami | Bitter | Sour | Salty |
| --- | --- | --- | --- | --- | --- | --- |
| SWE | 17 | 99 | 99 | 99 | 99 | 99 |
| SWE | 1 | 49 | 55 | 8 | 62 | 58 |
| SWE | 2 | 28 | 10 | 13 | 99 | 99 |
| SWE | 3 | 74 | 11 | 36 | 84 | 43 |
| SWE | 4 | 2 | 99 | 68 | 62 | 75 |
| SWE | 5 | 99 | 70 | 99 | 99 | 76 |
| SWE | 6 | 79 | 38 | 38 | 44 | 23 |
| SWE | 10 | 60 | 33 | 99 | 84 | 43 |
| SWE | 16 | 58 | 36 | 71 | 99 | 76 |
| SWE | 7 | 4 | 10 | 7 | 71 | 43 |
| SWE | 8 | 74 | 4 | 7 | 71 | 81 |
| SWE | 9 | 7 | 17 | 10 | 99 | 99 |
| SWE | 12 | 26 | 53 | 71 | 81 | 19 |
| SWE | 13 | 7 | 30 | 10 | 55 | 20 |
| SWE | 14 | 18 | 99 | 47 | 99 | 60 |
| SWE | 15 | 41 | 92 | 29 | 35 | 19 |
| SWE | 18 | 87 | 12 | 50 | 71 | 2 |
| SWE | 19 | 79 | 72 | 99 | 62 | 5 |
| SWE | 20 | 77 | 35 | 75 | 99 | 99 |
| SWE | 21 | 42 | 23 | 69 | 90 | 29 |
| SWE | 22 | 7 | 70 | 5 | 68 | 3 |
| SWE | 23 | 33 | 72 | 81 | 62 | 39 |
| SWE | 24 | 65 | 17 | 10 | 37 | 35 |
| SWE | 25 | 14 | 70 | 10 | 2 | 81 |
| SWE | 26 | 57 | 10 | 59 | 26 | 71 |
| SWE | 27 | 3 | 11 | 42 | 11 | 57 |
| SWE | 28 | 65 | 55 | 81 | 78 | 11 |
| SWE | 29 | 3 | 12 | 5 | 1 | 1 |
| Values≤10% |  | 6 | 2 | 8 | 1 | 4 |

